# Ketamine blocks morphine-induced conditioned place preference and anxiety-like behaviors in mice

**DOI:** 10.1101/2020.01.22.915728

**Authors:** Greer McKendrick, Hannah Garrett, Holly E. Jones, Dillon S. McDevitt, Sonakshi Sharma, Yuval Silberman, Nicholas M. Graziane

## Abstract

Patients suffering from opioid use disorder often relapse during periods of abstinence, which is posited to be caused by negative affective states that drive motivated behaviors. Here, we explored whether conditioning mice with morphine in a CPP training paradigm evoked anxiety-like behavior during morphine abstinence. To do this, mice were conditioned with morphine (10 mg/kg, i.p.) for five days. 24 h following conditioning, anxiety levels were tested by measuring time in the open arms of the elevated plus maze. The next day, mice were placed in the three compartment chamber to measure morphine-induced conditioned place preference (CPP). Our results show that following morphine conditioning, mice spent significantly less time in the open arm of the elevated plus maze and expressed robust morphine CPP on CPP test day.
Furthermore, we found that an acute treatment with (*R,S*)-ketamine (10 mg/kg, i.p.), a medication demonstrating promise for preventing anxiety-related phenotypes, 30 min. prior to testing on post conditioning day 1, increased time spent in the open arm of the elevated plus maze in saline- and morphine-conditioned mice. Additionally, we found that a second injection of ketamine 30 min. prior to CPP tests on post conditioning day 2 prevented morphine-induced CPP, which lasted for up to 28 d post conditioning. Furthermore, we found that conditioning mice with 10% (w/v) sucrose using an oral self-administration procedure did not evoke anxiety-like behavior, but elicited robust CPP, which was attenuated by ketamine treatment 30 min. prior to CPP tests. Overall, our results suggest that the ketamine-induced block of morphine CPP may not be attributed solely to alleviating negative affective states, but potentially through impaired memory of morphine-context associations.

## Introduction

The motivation to continually seek and obtain addictive substances during periods of abstinence or recovery is caused, in part, by the necessity to avoid aversive internal states (Solomon and Corbit, 1978). Evidence for this comes from patients with substance use disorders who self-report urges and intentions to take drugs to avoid drug-withdrawal symptoms (O’Brien, 1975;Baker et al., 2004;Wikler, 2013) or to cope with negative affect (Perkins and Grobe, 1992;Zinser et al., 1992;Wetter et al., 1994;Cooney et al., 1997;Conklin and Perkins, 2005;Fox et al., 2007). For example, abstinence from morphine, a highly addictive opioid, facilitates increases in anxiety (Gold et al., 1978; 1979), which is a potential factor in continued drug use (Martins et al., 2012).

In order to better understand the mechanisms mediating drug-craving and subsequent relapse, preclinical models have been developed whereby drug-seeking behaviors are monitored in drug-exposed rodents. In the conditioned place preference (CPP) paradigm, a drug is paired with a context during conditioning. This is followed by a test day whereby the time spent in the drug-paired context is measured. This behavioral paradigm is a form of Pavlovian learning whereby an injection of a drug (i.e., unconditioned stimulus) elicits a hedonic feeling of pleasure (i.e., unconditioned response), which, when paired with a context (neutral stimulus), invokes incentive value to the context (i.e., now a conditioned stimulus), thus driving a behavioral response to “seek” the context (conditioned response). This is similar to sign-tracking behaviors (Huston et al., 2013), which refer to a behavior that is directed toward a stimulus as a result of that stimulus becoming associated with a reward (Huys et al., 2014). Therefore, CPP provides a valuable tool used to understand how drugs of abuse become associated with environmental contexts, which is implicated in context-induced drug craving and relapse (O’Brien CP, 1986;O’Brien et al., 1992). We have found that five days of morphine (10 mg/kg) conditioning elicits robust morphine CPP (Graziane et al., 2016;McDevitt and Graziane, 2019). However, it is unclear whether this “drug context-seeking” behavior is mediated by negative affective states. Additionally, it is unclear whether a subanesthetic dose of ketamine, an anxiolytic agent (Engin et al., 2009b), blocks morphine-induced CPP by mitigating morphine-induced negative affective states.

Here, we attempt to investigate whether morphine conditioning in our CPP paradigm generates negative affect during morphine abstinence. Additionally, we investigate whether an acute, subanesthetic dose of (*R,S*)-ketamine prior to testing is sufficient to disrupt morphine-induced anxiety and/or morphine-induced CPP behaviors. Lastly, it has been shown that an acute administration of (*R,S*)-ketamine is sufficient to block the expression of morphine CPP (Suzuki et al., 2000). Here, we investigate whether this ketamine-induced block of morphine CPP, in our behavioral training paradigm, is mediated by the impairment of drug-context associations or by the attenuation of morphine-induced negative affective states.

## Methods

### Animals

All experiments were done in accordance with procedures approved by the Pennsylvania State University College of Medicine Institutional Animal Care and Use Committee. Male C57BL/6J mice aged 5-8 weeks were purchased from Jackson Labs (stock #000664) (Bar Harbor, ME), singly-housed, and maintained on a regular 12 hour light/dark cycle (lights on 07:00, lights off 19:00) with *ad libitum* food and water. Mice were singly housed for the following reasons. First, we have reliably developed morphine conditioned place preference (CPP) in singly-housed mice (Graziane et al., 2016;McDevitt and Graziane, 2019). Second, evidence suggests that socially isolated rodents are more vulnerable to developing drug-context associations (Whitaker et al., 2013). In humans, social isolation increases vulnerability to substance use disorders (Newcomb and Bentler, 1988;Sinha, 2008), which often are accompanied by the development of drug-context associations (O’Brien CP, 1986;O’Brien et al., 1992;Xue et al., 2012). Therefore, our studies are designed to model this patient population.

### Drugs

(−)-morphine sulfate pentahydrate was provided by the National Institute on Drug Abuse Drug Supply Program. Ketamine hydrochloride (racemic mixture of 50% *R*-ketamine and *S*-ketamine) (Dechra Pharmaceuticals, Northwich, United Kingdom) was purchased from the Comparative Medicine Department at the Pennsylvania State University College of Medicine.

### Non-Contingent Conditioned Place Preference

Conditioned place preference (CPP) chambers (Med Associates) were located in the mouse housing room and consisted of three distinct compartments separated by manual guillotine-style doors. Each compartment had distinct contextual characteristics: the middle (neutral) compartment (7.2 cm × 12.7 cm × 12.7 cm) had grey walls and grey plastic floor, while the choice compartments (16.8 cm × 12.7 cm × 12.7 cm, each) had either white walls and stainless steel mesh floor or black walls and stainless steel grid floor. All compartments were illuminated with a dim light during use. Immediately following use the entire preference chamber was cleaned thoroughly with a scent-free soap solution. Mouse locations, activity counts, and time spent in each compartment were collected via automated data-collection software (Med Associates) via infrared photobeam strips lining each compartment. Morphine administration was verified with the Straub tail response and enhanced locomotor activity (Bilbey et al., 1960;Graziane et al., 2016;McDevitt and Graziane, 2019).

Habituation. Mice were placed in the center compartment with free access to all three compartments for 20 min once a day for two days. Time spent (seconds) in each compartment was recorded.

Conditioning. 24 h after habituation, mice received 5 d conditioning training. Morphine-paired compartments were assigned based on the least preferred side (a biased approach) (Tzschentke, 2007) calculated by averaging time spent in each compartment over the 2 habituation days. Similar to conditioning studies with alcohol (Gremel et al., 2006), we find that C57BL/6J mice will reliably develop morphine CPP using a biased approach. During conditioning, mice received an injection of saline and were placed into the most preferred compartment for 40 min. 6 h later, mice received an injection of saline (control group) or morphine (10 mg/kg, i.p.) and were placed into their least preferred compartment for 40 min. (Koo et al., 2014;Graziane et al., 2016).

Post conditioning. 48 h or 28 d after the last conditioning day, mice were placed in the 3-compartment chamber and allowed to move freely for 20 min. Our post-conditioning took place at a time point corresponding to 3 h prior to drug conditioning (e.g., morphine conditioning took place at 3 P.M., post-conditioning tests took place 2 or 28 days later at 12 P.M.). CPP scores were calculated as time spent in the drug-paired side minus the average time spent on the same side during preconditioning (Bohn et al., 2003). Activity counts are defined as any beam break within a current zone. This is inclusive of grooming, rearing, and lateral movements. Mice were treated with 0.9% saline (0.1 ml, i.p.) or with (*R,S*)-ketamine (10 mg/kg, i.p.) 30 min. prior to the first CPP test. The dose of ketamine was selected based on preclinical data demonstrating that a 10 mg/kg dose of ketamine produces a maximal effect on morphine CPP (Suzuki et al., 2000) and produces plasma concentrations associated with subanesthetic ketamine doses capable of eliciting antidepressant effects in mice and in humans (Zarate et al., 2012;Zanos et al., 2016).

### Sucrose Oral Self-Administration Conditioned Place Preference

Habituation. Mice were placed in the center compartment with free access to all three compartments for 20 min. once a day for two days. Time spent (seconds) in each compartment was recorded.

Conditioning. Drinking bottles were created as described in Freet et al., 2013 (Freet et al., 2013). Briefly, we modified 10 mL serological pipettes by tapering both ends, placing a stainless-steel sipper tube (Ancare; OT-300) in one end and a silicon stopper (Fisher Scientific; 09-704-1D) in the other. Bottles were inserted into plastic holders that were then placed directly into CPP chambers (for chamber description, see Non-Contingent Conditioned Place Preference), where they were positioned so that the sipper was ~5 cm above the chamber floor. Pennsylvania State University Fabrication shop constructed plexiglass tops that were placed along the top of the 3-compartment apparatus and allowed for plastic bottle holders to be placed into chambers. Oral self-administration was recorded as the mL prior and following all sessions. Similar to the i.p. CPP methodology, we utilized a biased approach in which the 10% sucrose (w/v) solution was placed in the least-preferred context. 24 h after habituation, mice underwent two 14 h overnight sessions (separated by 24 h), confined to the least preferred chamber on the first night (ON1) with access to water (control groups) or a 10% sucrose solution and confined to the most preferred side on the second night (ON2) with access to water. Mice then received 5 days of conditioning (C1-C5), where morning sessions consisted of 40 min. in the most-preferred context with access to water. 6 h later, afternoon sessions consisted of 40 min. in the least preferred context with access to water (control groups) or 10% sucrose solution.

Post conditioning. 48 h or 21 d after the last conditioning day, mice were placed in the 3-compartment chamber and allowed to move freely for 20 min. Our post-conditioning took place at a time point corresponding to 3 h prior to drug conditioning (e.g., sucrose conditioning took place at 3 P.M., post-conditioning tests took place 2 or 21 days later at 12 P.M.). No bottles were present in the chambers on preference tests. CPP scores were calculated as time spent in the least preferred side on test day minus the average time spent on the same side during preconditioning (Bohn et al., 2003). Mice treated with (R,S)-ketamine (10 mg/kg, i.p.) (water+ketamine and sucrose+ketamine groups) received injections 30 min. prior to the first CPP test on post conditioning day 2.

### Elevated Plus Maze

The elevated-plus maze, a well-established method to measure anxiety in rodents, was implemented to measure anxiety-like behavior (Pellow et al., 1985b;Handley and McBlane, 1993;Dawson and Tricklebank, 1995). The elevated-plus maze for mice (Stoelting, Item #60140) was raised approximately 50 cm from the ground. The floor of the elevated portion of the maze was gray. Two opposite arms (35 × 5 cm each) of the maze were enclosed by a 15 cm high wall and the remaining two arms were “open.” A center space (5 cm^2^) between these four arms was also not enclosed. The elevated portion of the apparatus was cleaned thoroughly with a scent-free soap solution after each trial. Behavioral tests were performed in the animal housing room under ambient light of the light cycle.

24 h after the last conditioning day in the CPP apparatus, mice were placed in the center space facing the open arm and allowed to explore the apparatus for 5 minutes prior to being placed back into their home cage (Grisel et al., 2008). Each trial was video recorded using a GoPro camera (Hero7 white) and analyzed by researchers blinded to treatment condition of the mice. Time in the open arm was measured when the body of the mouse cleared the center space. Mice were treated with 0.9% saline (0.1 ml, i.p.) or ketamine (10 mg/kg, i.p.) 30 min. prior to the elevated plus maze test.

### Statistical Analysis

Statistical significance was assessed in GraphPad Prism software using a Student’s t-test, one- or two-way ANOVA with Bonferroni’s correction for multiple comparisons as specified. F values for two-way ANOVA statistical comparisons represent interactions between variables unless stated otherwise. Two-tailed tests were performed for Student’s t-test. For correlation analysis, the Pearson’s correlation coefficient, and subsequent linear regression, were determined. P<0.05 was considered to indicate a statistically significant difference.

## Results

### Morphine conditioning elicits anxiety-like behaviors during morphine abstinence

Repeated exposure to morphine increases levels of anxiety both in humans and in animal models of substance use disorders (Gold et al., 1978;1979;Becker et al., 2017). Additionally, it is posited that relapse to opioids in abstinent patients is caused by negative affective states, thus driving drug-seeking behaviors (Solomon and Corbit, 1978;Koob and Le Moal, 2008;Evans and Cahill, 2016). In an attempt to provide evidence that morphine-induced CPP, using our training paradigm, is mediated, in part, by negative affective states, 24 h following the last morphine conditioning session (**Fig. 1A**), we measured anxiety-like behavior using the elevated plus maze (EPM) (Pellow et al., 1985a). We found that morphine-treated mice, who showed robust locomotor sensitization by conditioning day 5 (**Fig. 1B**), expressed a significant decrease in the percent time spent in the open arm of the EPM compared to saline-treated controls (t_(38)_=3.35, p=0.002, Student’s t-test) (**Fig. 1C**). To correlate anxiety levels with CPP scores, mice underwent CPP tests 24 h following EPM tests (**Fig. 1A**). We found that 5 d morphine conditioning elicited significant increases in place preference for the drug-paired compartment (t_(38)_=5.61, p<0.0001, Student’s t-test) (**Fig. 1D**). However, we found no correlation between anxiety-like behaviors and CPP score in morphine-conditioned mice (Pearson’s correlation coefficient = −0.162; simple linear regression: F_(1,15)_=0.404, p=0.53, R^2^ =0.03) or in saline-conditioned, control mice (Pearson’s correlation coefficient = −0.095; simple linear regression: F_(1,21)_=0.191, p=0.67, R^2^=0.01) (**Figs. 1E and F**). Overall, these results suggest that morphine conditioning in a CPP paradigm is sufficient to facilitate anxiety-like behaviors during short-term abstinence, but that the animal’s anxiety-like behavior is not correlated with the amount of time spent in the morphine-paired compartment on CPP test day.

**Figure 1.**
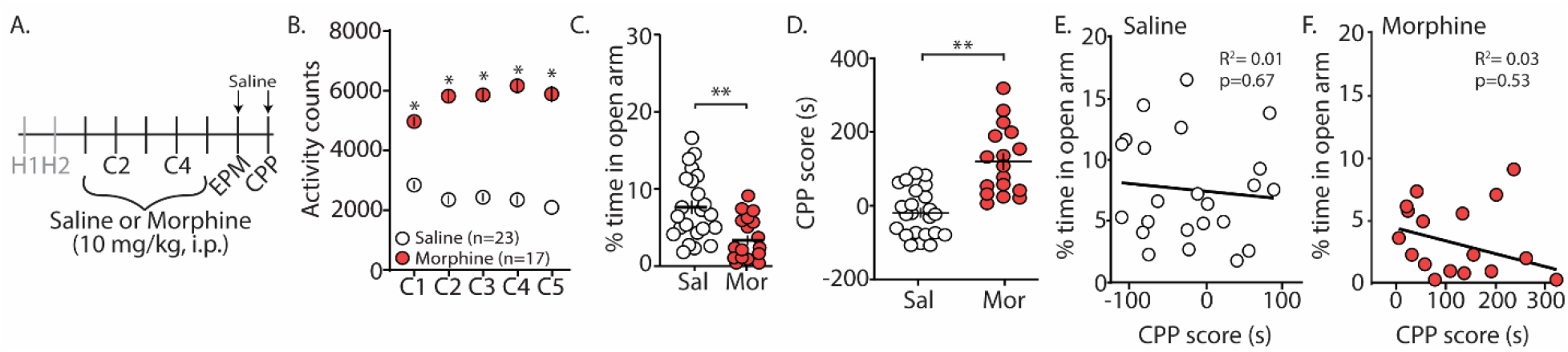
Morphine conditioning in a CPP paradigm elicits anxiety-like behaviors during 24 h abstinence. (A) Time line and drug regimen of the behavioral procedure. Animals underwent two days of habituation (H), followed by five days of saline or morphine (10 mg/kg, i.p.) conditioning (C), before being subjected to tests measuring anxiety-like behaviors using an elevated plus maze (EPM) 24 h post conditioning. 24 h post EPM tests, CPP tests were performed. Animals were injected with saline 30 min. prior to EPM and CPP tests. (B) Summary showing that morphine conditioning over 5 days produces robust locomotor sensitization (F_(4, 152)_= 17.1, p<0.0001, two-way repeated measures ANOVA, Bonferroni post hoc test). (C) Summary showing that morphine (Mor)-conditioned mice spent significantly less time in the open arms of the elevated plus maze compared to saline (Sal)-conditioned mice 24 h following the last conditioning day (t_(38)_=3.35, p=0.002, Student’s t-test). (D) Summary showing that morphine conditioning produced reliable CPP (t_(38)_=5.61, p<0.0001, Student’s t-test). (E) Correlation of the % time in the open arm of the elevated plus maze and CPP score in saline- or (F) morphine-conditioned mice. *p<0.05, **p<0.01.

### Ketamine blocks morphine-induced anxiety-like behaviors and morphine CPP

Evidence suggests that (*R,S*)-ketamine, a noncompetitive NMDA receptor antagonist (Lodge et al., 1982;Kohrs and Durieux, 1998), is an effective treatment for anxiety and substance use disorders (Krupitsky et al., 2002a;Ivan Ezquerra-Romano et al., 2018;Taylor et al., 2018). Because of this, we investigated whether an acute injection of (*R,S*)-ketamine (30 min. prior to EPM and CPP testing) would be sufficient to block morphine-induced anxiety-like behaviors and/or morphine-induced CPP (**Fig. 2A**). Following conditioning with morphine, which produced robust locomotor sensitization (**Fig. 2B**), we found that the first (*R,S*)-ketamine injection prior to the EPM test on post-conditioning day 1 (PC1) significantly increased the percent time in the open arms of the EPM (F_(3, 52)_=22.2, p<0.0001, one-way ANOVA, Bonferroni post hoc test) (**Fig. 2C**). Additionally, we found that a second (*R,S*)-ketamine injection prior to CPP tests on post-conditioning day 2 (PC2) was sufficient to prevent morphine-induced CPP (F_(3, 52)_=14.04, p<0.0001, one-way ANOVA, Bonferroni post hoc test) (**Fig. 2D**), which was likely not attributed to ketamine-induced changes in locomotor activity (F_(3,52)_=0.447, p=0.72, two-way repeated measures ANOVA) (**Fig. 2E**).

**Figure 2.**
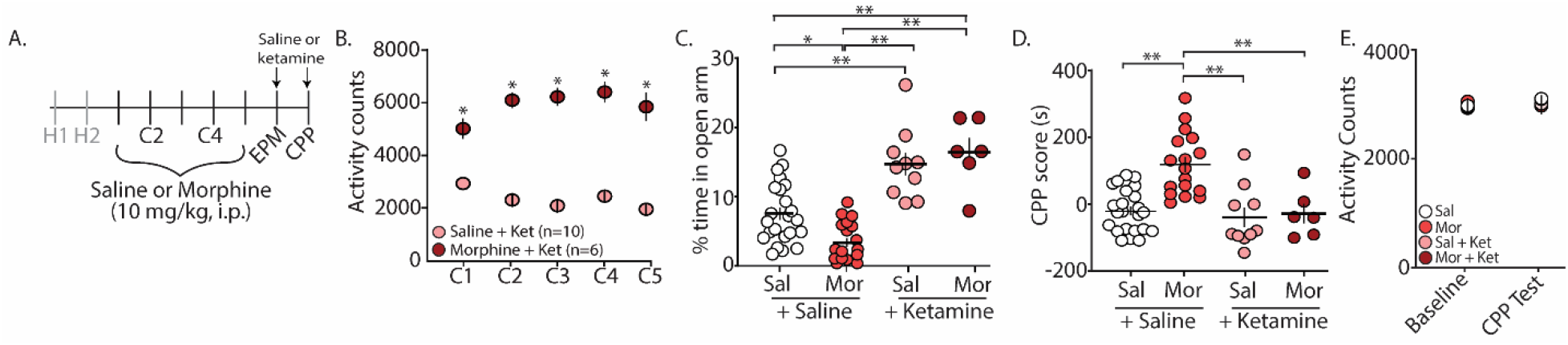
Acute (*R,S*)-ketamine injection produces anxiolytic-like behaviors in mice 24 h after conditioning and blocks morphine-induced CPP. (A) Time line and drug regimen of the behavioral procedure. Saline or (*R,S*)-ketamine (10 mg/kg, i.p.) was injected 30 min. prior to elevated plus maze (EPM) test with the second injection taking place 30 min. prior to the first conditioned place preference (CPP) test. (B) Summary showing that morphine conditioning over 5 days (C1-C5) produces robust locomotor sensitization (F_(4, 56)_= 12.55, p<0.0001, two-way repeated measures ANOVA, Bonferroni post hoc test). (C) Summary showing that (*R,S*)-ketamine significantly increased the time spent in the open arms of the elevated plus maze in both saline (Sal)- and morphine (Mor)-conditioned mice (F_(3, 52)_=22.2, p<0.0001, one-way ANOVA, Bonferroni post hoc test) (animals not receiving (*R,S*)-ketamine are the same data as shown in Fig. 1C). (D) Summary showing that morphine produced reliable CPP at post conditioning day 2, which was blocked by (*R,S*)-ketamine injected 30 min prior to testing (F_(3, 52)_=14.04, p<0.0001, one-way ANOVA, Bonferroni post hoc test) (saline and morphine groups are the same animals as shown in Fig. 1D). (E) Summary showing the activity counts in the CPP chamber during habituation (baseline) and during the CPP test in saline (Sal)- or morphine (Mor)-conditioned mice treated with saline or (*R,S*)-ketamine 30 min prior to testing (F_(3,52)_=0.447, p=0.72, two-way repeated measures ANOVA). *p<0.05, **p<0.01.

### Acute ketamine treatment blocks the long-term expression of morphine CPP

We have previously shown that morphine-induced CPP, using the paradigm described in this study, is sufficient to elicit long-lasting CPP for up to 28 d post conditioning (Graziane et al., 2016). Because of this, we tested whether ketamine administration during early abstinence was sufficient to block the prolonged expression of morphine-induced CPP (**Fig. 3A**). We found that two injections of (*R,S*)-ketamine, one on post conditioning day 1 (prior to elevated arm maze tests) and the second on post conditioning day 2 (prior to CPP tests), was sufficient to prevent the prolonged expression of morphine-induced CPP on PC28 (column factor: F_(3, 38)_=10.25, p<0.0001, two-way repeated measures ANOVA, Bonferroni post hoc test) (**Fig. 3B**).

**Figure 3.**
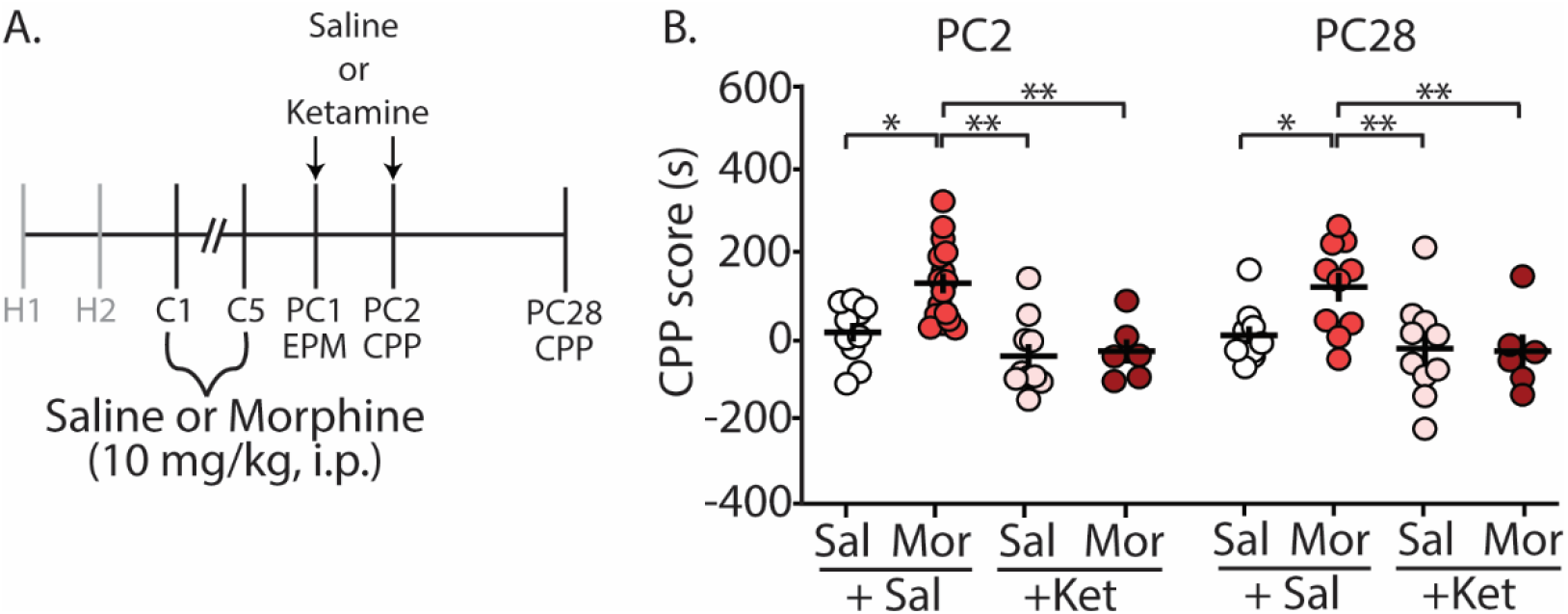
(*R,S*)-ketamine administration during early abstinence is sufficient to prevent the prolonged retention of morphine-induced CPP at post conditioning day 28. (A) Time line and drug regimen of the behavioral procedure. (*R,S*)-ketamine (10 mg/kg, i.p.) was injected 30 min. prior to the EPM test on post-conditioning day 1 (PC1) and again on the first CPP test on post conditioning day 2 (PC2) (i.e., each mouse received a ketamine injection before the EPM test and a second ketamine injection the next day prior to the CPP test). The second CPP test was run on PC28. (B) Summary showing that morphine produced reliable CPP 28 d post conditioning, which was blocked by (*R,S*)-ketamine (column factor: F_(3, 38)_=10.25, p<0.0001, two-way repeated measures ANOVA, Bonferroni post hoc test) (PC2 data is the same data shown in Fig. 2D). Abbrev.: EPM=elevated plus maze; CPP=conditioned place preference. *p<0.05, **p<0.01.

### Acute ketamine treatment prevents the expression of sucrose CPP

To further investigate whether the ketamine block of morphine CPP is through potential memory impairment and/or anxiolytic effects, we evaluated the effect of ketamine on the CPP of a natural reward (i.e., sucrose). We rationalized that if ketamine blocks morphine CPP by specifically alleviating negative affective states, without impairing memory of drug-context associations, then ketamine would be ineffective at blocking sucrose CPP, a natural reward, which does not evoke anxiety-like behaviors (**Fig. 4C**). To test this, we conditioned mice over 7 days (**Fig. 4A**) to orally self-administer water (controls) or sucrose in the least preferred compartment of the CPP chamber (see Methods for conditioning paradigm). Mice conditioned with sucrose drank significantly more than mice conditioned with water over all conditioning days (F_(15, 175)_= 462.1, p<0.0001, two-way repeated measures ANOVA, Bonferroni post hoc test) (**Fig. 4B**). The water consumed in the most preferred chamber during conditioning days 1-5 did not differ between groups (F_(12, 140)_=0.596, p=0.843, two-way repeated measures ANOVA) (**Supplementary Figure 1**). On post-conditioning day 1 (PC1), anxiety-like behavior was measured using the EPM. We found that the percent time in the open arm of the EPM in sucrose-conditioned mice was not significantly different from mice conditioned with water (t_(17)_=0.184, p=0.856, Student’s t-test) (**Fig. 4C**) suggesting that sucrose exposure did not elicit anxiety-like behaviors during short-term abstinence. 24 h later, on post-conditioning day 2 (PC2), water- and sucrose-conditioned mice underwent a CPP test 30 min. after receiving an acute injection of (*R,S*)-ketamine (10 mg/kg, i.p.). Our data show that (*R,S*)-ketamine attenuated sucrose-induced CPP on PC2 (F_(3, 35)_=6.31, p=0.0015, one-way ANOVA, Bonferroni post hoc test) (**Fig. 4D**) and this ketamine-induced attenuation of sucrose CPP persisted to abstinence day 21 (F_(3, 32)_=5.51, p=0.004, one-way ANOVA, Bonferroni post hoc test) (**Supplementary Figure 2**).

**Figure 4.**
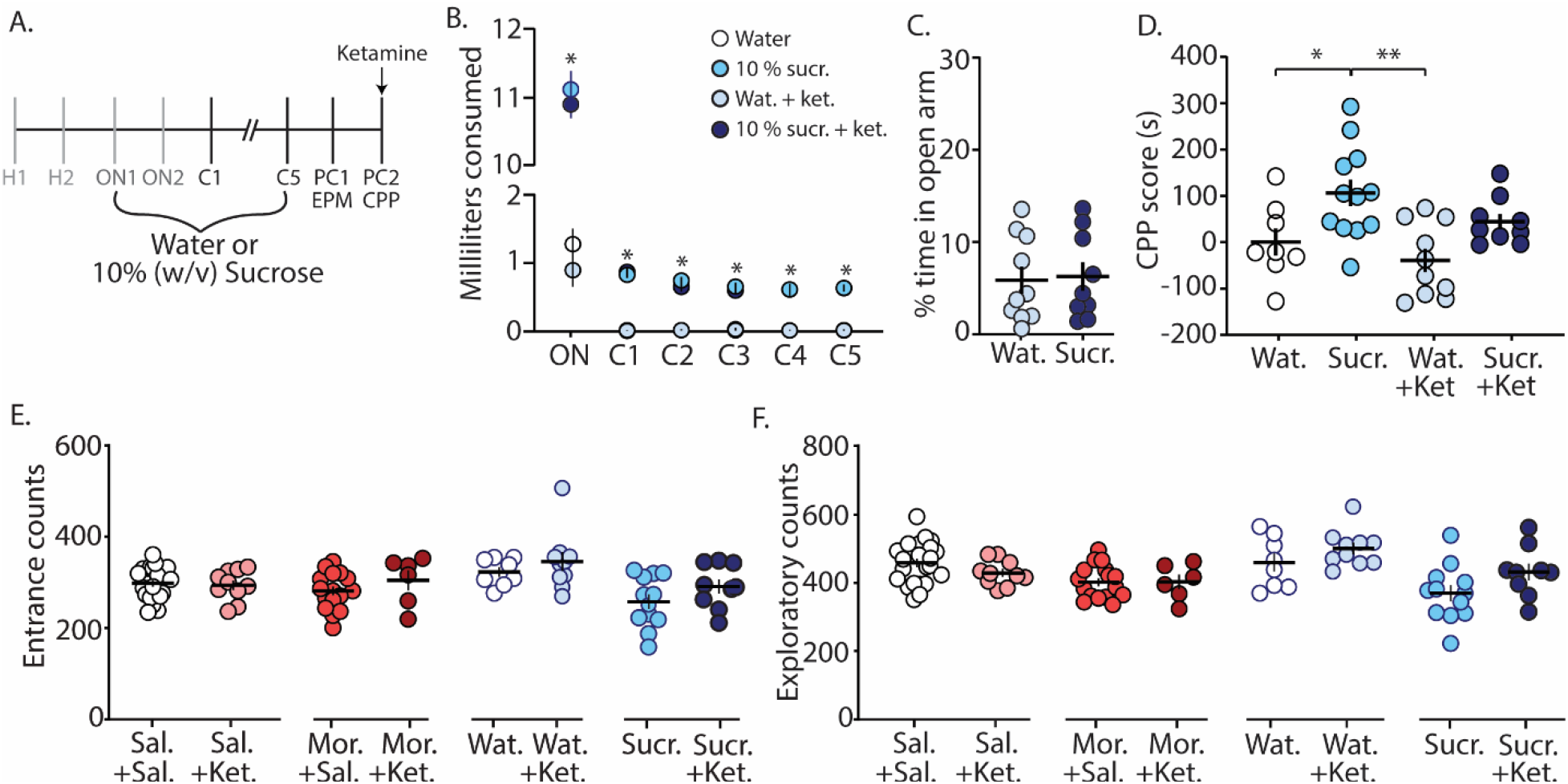
Ketamine administration attenuates sucrose-induced conditioned place preference. (A) Time line and sucrose regimen of the behavioral procedure. Following sucrose oral self-administration in the three compartment apparatus, mice underwent EPM testing on post-conditioning day 1 (PC1). 24 h later, mice received no injection or (*R,S*)-ketamine (10 mg/kg, i.p.) 30 min. prior to the conditioned place preference (CPP) test on post-conditioning day 2 (PC2). (B) Summary showing the milliliters of water or sucrose consumed for each training session in the least preferred chamber. Groups conditioned with sucrose (i.e., sucrose (sucr.) and sucrose+ketamine (sucr.+ket.) groups) drank significantly more than groups conditioned with water (i.e., water (Wat.) and water+ketamine (Wat.+Ket.) groups) (F_(15, 175)_= 462.1, p<0.0001, two-way repeated measures ANOVA, Bonferroni post hoc test). (C) Summary showing that conditioning with sucrose had no effect on anxiety-like behaviors as both water- and sucrose-conditioned mice displayed similar % time in the open arm of the EPM (t_(17)_=0.184, p=0.856, Student’s t-test). (D) Summary showing that oral self-administration of sucrose produced CPP at PC2, which was blocked by (*R,S*)-ketamine treatment (F_(3, 35)_=6.31, p=0.0015, one-way ANOVA, Bonferroni post hoc test). (E) Summary showing that ketamine injections 30 min. prior to CPP test did not impact entrance counts in the CPP apparatus (Sal.+Sal. vs. Sal.+Ket.: t_(31)_=0.295, p=0.770; Mor.+Sal. vs. Mor.+Ket.: t_(21)_=1.13, p=0.272; Wat.+Sal. vs. Wat.+Ket.: t_(16)_=0.874, p=0.395; Sucr..+Sal. vs. Sucr.+Ket.: t_(19)_=1.43, p=0.168, Student’s t-test). (F) Summary showing that ketamine injections 30 min. prior to CPP test did not impact exploratory counts in the CPP apparatus (Sal.+Sal. vs. Sal.+Ket.: t_(31)_=1.42, p=0.166; Mor.+Sal. vs. Mor.+Ket.: t_(21)_=0.045, p=0.964; Wat.+Sal. vs. Wat.+Ket.: t_(16)_=1.26, p=0.226; Sucr..+Sal. vs. Sucr.+Ket.: t_(19)_=1.80, p=0.088, Student’s t-test). *p<0.05, **p<0.01.

Lastly, we investigated whether the ketamine block of morphine-induced anxiety-like behavior and morphine-induced CPP was potentially attributed to ketamine-induced behavioral disinhibition, leading the animal to explore more. To do this, we monitored entrance counts and exploratory counts in the CPP chamber on test day. We found that there was no significant difference in entrance or exploratory counts in the CPP chamber when comparisons were made between saline versus ketamine injected mice undergoing the same treatment during conditioning (**Figs. 4E and F**). These results suggest that the effects of ketamine on morphine-driven behaviors is unlikely mediated by behavioral disinhibition.

## Discussion

Our results show that the percent time spent in the open arms of the elevated plus maze is decreased in animals conditioned with morphine. Additionally, we show that an acute injection of (*R,S*)-ketamine 30 min prior to the elevated plus maze and CPP tests is sufficient to block morphine-induced anxiety-like behaviors and morphine-induced CPP (post-conditioning day 2 through post-conditioning day 28), as well as attenuates sucrose-induced CPP (post-conditioning day 2 through post-conditioning day 21). We further find that ketamine, at least in the dose tested here, does not alter behavioral disinhibition in either morphine-CPP or sucrose-CPP mice. Together these findings indicate that ketamine may inhibit morphine CPP behaviors, at least in part, via reductions in withdrawal-induced anxiety-like behaviors. Our data do not, however, rule out the possibility that ketamine-induced effects on morphine CPP may also be mediated in part by impairing memory of morphine-context associations.

### Anxiety-like behaviors during morphine abstinence

Morphine possesses anxiolytic-like properties during initial exposure (Koks et al., 1999;Sasaki et al., 2002;Shin et al., 2003). However, during opioid abstinence, symptoms of anxiety (Gold et al., 1978;1979;Li et al., 2009;Shi et al., 2009) or anxiety-like behaviors are observed (Cabral et al., 2009;Becker et al., 2017). Here, we show that 24 h following repeated morphine injections (once a day for 5 days), mice display anxiety-like behaviors in the elevated plus maze (**Fig. 1C**). These results are similar to previous studies showing escalating doses of morphine over a 6 day period induce anxiety-like behaviors in the marble burying task (Becker et al., 2017). Additionally, our observed morphine-induced anxiety-like behavior is timed with anxiogenic neurobiological responses that occur during acute opioid abstinence including, increases in norepinephrine release in the extended amygdala (Fuentealba et al., 2000;Aston-Jones and Harris, 2004), norepinephrine-induced modulation of the extended amygdala (Aston-Jones et al., 1999;Delfs et al., 2000;Smith and Aston-Jones, 2008), activation of the amygdalar corticotrophin-releasing factor (CRF) system (Heinrichs et al., 1995;Maj et al., 2003), and decreases in dopamine transmission (Diana et al., 1995). However, the observed morphine-induced anxiety-like behavior may be dependent upon morphine exposure as it has been shown that morphine does not elicit anxiety-like behaviors following three morphine injections (10 mg/kg) occurring every other day (Benturquia et al., 2007). This may be related to neurobiological mechanisms associated with different drug exposure regimens. We have previously shown that morphine exposure significantly increases the expression of silent synapses, excitatory glutamatergic synapses that express functional NMDA receptors, but lack functional AMPA receptors (Hanse et al., 2013), in the nucleus accumbens shell. We found that this increase in silent synapse expression is observed 24 h after the last of five morphine injections (once a day for five days), but not 24 h after the last of three morphine injections (once a day for three days) (Graziane et al., 2016;Hearing et al., 2018;McDevitt and Graziane, 2018). Future experiments will be required to test whether this morphine-induced change in the nucleus accumbens shell regulates morphine-induced anxiety-like behaviors.

The observed anxiety-like behaviors following morphine conditioning in a three chamber apparatus (**Fig. 1F**) may suggest that animals seek the drug-paired chamber as a consequence of negative reinforcement to alleviate aversive affective states facilitated by opioid abstinence. Importantly, our injection regimen of morphine 10 mg/kg once a day for 5 consecutive days does not induce signs of somatic withdrawal in mice including jumping, wet dog shakes, teeth chattering, rearing, tremor, diarrhea, or mastication (Gallego et al., 2010). This coincides with the lack of observed somatic withdrawal symptoms following a more prolonged injection regimen of 5 daily morphine (10 mg/kg, i.p.) injections over 4 weeks (Robinson and Kolb, 1999). Although more studies are required, it is plausible that specific opioid dosing regimens may be implemented in a preclinical setting in order to separate opioid-induced negative affective states (e.g., anxiety) from confounds induced by somatic signs of opioid withdrawal, which are ineffective at reinstating opioid seeking or morphine CPP in opioid dependent rodents (Shaham et al., 1996;Lu et al., 2005) as well as in humans (Miller et al., 1979). Separating opioid-induced negative affective states (e.g., anxiety) from confounds induced by somatic signs of opioid withdrawal is not a new idea and has been demonstrated previously with doses of naloxone (used to precipitate opioid withdrawal) that were sub-threshold for somatic signs of opioid withdrawal (Gracy et al., 2001).

Based on our results, it would be expected that facilitating a negative affective state during morphine abstinence would enhance the expression of morphine CPP. However, evidence suggests that this is not the case, as forced swim stress, which would be expected to elicit a strong negative affective state, immediately prior to CPP testing in morphine-conditioned animals has either no effect on morphine CPP (Attarzadeh-Yazdi et al., 2013) or significantly decreases morphine CPP (Haghparast et al., 2014). Additionally, corticosterone administration, which is expected to facilitate depression-like behaviors (Gregus et al., 2005), prior to CPP tests has no effect on morphine CPP (Attarzadeh-Yazdi et al., 2013). These results are surprising especially considering the robust effect of stressful stimuli in reinstating morphine CPP in extinguished rodents (Ribeiro Do Couto et al., 2006;Wang et al., 2006;Karimi et al., 2014). It is possible that morphine CPP tested during abstinence (e.g., Attarzadeh-Yazdi et. al., 2013) reaches a ceiling effect, making it unlikely that exposure to a stressor (e.g., forced swim) will enhance the CPP score (i.e., occlusion). It is also possible that the stressor elicits a decreased locomotor state potentially resulting in reduced morphine CPP (e.g., Haghparast et al., 2014).

### Ketamine’s effects on anxiety-like behaviors

Ketamine has recently been shown to be a potential effective treatment for anxiety disorders (Glue et al., 2018;Shadli et al., 2018;Taylor et al., 2018). In humans, ketamine displays a biphasic dose effect on anxiety, with low doses decreasing anxiety and higher doses increasing anxiety (Jansen, 1989;Krystal et al., 1994). Likewise, in rodents, ketamine induces anxiolytic-like behaviors (Engin et al., 2009a;Zhang et al., 2015;Fraga et al., 2018) as well as anxiogenic-like phenotypes likely dependent upon the dose, temporal relationship between ketamine injection and test onset, and rodent species (Silvestre et al., 1997;da Silva et al., 2010). Here, we demonstrate that in C57BL/6J mice, acute injection of ketamine at 10 mg/kg, i.p. 30 min prior to testing is sufficient to block morphine-induced anxiety-like behaviors during a 24 h abstinence time period (**Fig. 2C**). Additionally, we find that ketamine significantly increases percent time in the open arm of the elevated plus maze in mice conditioned with saline. This significant change observed in saline conditioned animals suggests that ketamine, at the dose and temporal relationship of ketamine injection and test onset, is sufficient to overcome baseline anxiety-like behaviors in animals exposed to a novel environment (i.e., elevated plus maze).

Despite the evidence suggesting that the antagonistic effects of ketamine on NMDA receptors in the bed nucleus of the stria terminalis attenuate negative affective states (Louderback et al., 2013), the mechanisms mediating the observed anxiolytic-like effects are unknown. In addition to acting as a non-competitive antagonist to NMDA receptors in the extended amygdala, evidence suggests that ketamine interacts with hyperpolarization-activated cyclic nucleotide-gated (HCN) channels as well as dopamine, serotonin, sigma, opioid, and cholinergic receptors (Scheller et al., 1996;Cai et al., 1997;Kubota et al., 1999;Lydic and Baghdoyan, 2002;Wang et al., 2012;Zanos et al., 2018). Additionally, ketamine metabolites are biologically active as antagonists to NMDA receptors (Ebert et al., 1997) and α7 nicotinic acetylcholine receptors (Moaddel et al., 2013), while also possessing agonistic activity for α-amino-3-hydroxy-5-methyl-4-isoxazolepropionic acid (AMPA) receptors (Zanos et al., 2016;Tyler et al., 2017). Because of the undiscriminating activity of ketamine and its metabolites, it has been difficult to pinpoint how ketamine influences anxiety states both in humans and in preclinical models.

### Ketamine’s effects on morphine-induced conditioned place preference

Using a paradigm known to induce robust CPP for up to 28 d post conditioning (Graziane et al., 2016), we show that an acute injection of ketamine 30 min prior to the CPP test on abstinence day 2 is sufficient to block morphine-induced CPP. These results are not likely caused by changes in locomotor activity as activity counts during habituation (baseline) were not significantly different from activity counts measured following ketamine administration (**Fig. 2E**). Our results are in line with previous publications demonstrating that ketamine blocks morphine-induced CPP in mice (Suzuki et al., 2000). However, the effects on locomotor activity are conflicting. Whereas, our results and those from previous publications show that ketamine does not influence locomotor activity (Lindholm et al., 2012), others have found that locomotor activity is increased (Filibeck and Castellano, 1980) or decreased following ketamine administration (Akillioglu et al., 2012). These discrepancies are likely due to the temporal relationship between ketamine treatment and test onset. Here, we performed our tests 30 min following ketamine injection similar to previous studies (Lindholm et al., 2012), while tests performed 5 min or 15 min following ketamine administration appear to increase or decrease locomotor activity, respectively (Filibeck and Castellano, 1980;Akillioglu et al., 2012). The half-life of ketamine is ~13-25 min. in mice following i.p. administration (Maxwell et al., 2006;Zanos et al., 2016;Ganguly et al., 2018). Therefore, it is possible that the locomotor effects observed are due to ketamine action prior to metabolism, while the effects on negative affect are potentially attributed to ketamine metabolites including hydroxynorketamine (Li et al., 2015;Zanos et al., 2016). This hypothesis will need to be tested in future experiments. Moreover, our results are based on using a fixed dose of ketamine at 10 mg/kg, thus preventing dose-response observations. Future investigations are required to test how varying ketamine doses may influence morphine-induced conditioned place preference as well as morphine-induced anxiety-like behaviors.

Based on our findings that ketamine elicited anxiolytic-like behaviors following an acute injection, it is possible that the acute administration of ketamine was sufficient to prevent a negative affective state during 24 h morphine abstinence, thus facilitating the lack of motivation to seek a context paired with a drug reward (i.e., morphine-induced CPP). It is also plausible that the block of morphine-induced CPP by ketamine may be mediated by its effects on cognition and memory, thus blocking the recall of morphine-context associations (Ghoneim et al., 1985;Newcomer et al., 1999;Morgan et al., 2004) (Malhotra et al., 1996;Pfenninger et al., 2002). Evidence suggests that ketamine-induced deficits in cognitive functioning and memory occur during the consolidation or, as shown in rodents, reconsolidation (Zhai et al., 2008) of information, rather than the retrieval of already learned associations (Honey et al., 2005). Furthermore, it has been shown in rodent models that the memory impairing effects of ketamine are not attributed to its effects on memory retrieval (Goulart et al., 2010). Therefore, an acute injection of ketamine prior to CPP tests is not likely to influence already encoded morphine-context associations. However, we found that ketamine was effective at attenuating sucrose-induced CPP, despite the lack of anxiety-like behavior induced by sucrose conditioning (**Figs. 4C and D**). Therefore, these data suggest that ketamine is able to interfere with memory associated with Pavlovian learning when administered prior to retrieval of already learned associations. We acknowledge that our data does not unequivocally demonstrate that the ketamine-induced block of morphine CPP is solely mediated by impairing already learned associations. Therefore, future studies are required to test whether blocking only morphine-induced negative affective states are sufficient to prevent morphine CPP.

Lastly, our data suggest that the effects of ketamine on morphine-induced anxiety-like behavior and on morphine CPP is not likely a result of ketamine-induced behavioral disinhibition, which would be expected to increase exploratory behaviors. We found that ketamine had no effect on entrance counts or exploratory behaviors in the CPP apparatus (**Fig. 4E and F**).

Overall, our data suggest that ketamine may influence morphine CPP by altering negative affective states as well as by altering memory of learned associations. However, this does not rule out that ketamine’s effects on morphine-induced CPP may be mediated by other mechanisms of action as ketamine has proven effective for treating pain (Weisman, 1971;Laskowski et al., 2011;Jonkman et al., 2017), depression (Khorramzadeh and Lotfy, 1973;Sofia and Harakal, 1975), and inflammation (Roytblat et al., 1998;Beilin et al., 2007;Loix et al., 2011).

### Ketamine as a treatment option for substance use disorders

There is growing clinical and preclinical evidence that ketamine may be a potential treatment option for substance use disorders (Ivan Ezquerra-Romano et al., 2018;Jones et al., 2018). Through the use of Ketamine Assisted Psychotherapy (KAP) (Ivan Ezquerra-Romano et al., 2018), alcohol-dependent patients (Krupitsky and Grinenko, 1997;Kolp et al., 2006), heroin-dependent patients (Krupitsky et al., 2002b;Krupitsky et al., 2007), and cocaine-dependent patients (Dakwar et al., 2017) showed greater rates of abstinence and reductions in drug craving. These results have been echoed in preclinical models of substance use disorders as acute ketamine injections significantly attenuate alcohol self-administration (Sabino et al., 2013) and prevent the reconsolidation of morphine-induced CPP (Zhai et al., 2008). Here, we discovered a novel and unexpected loss of long-term expression of morphine-induced CPP in animals injected with (*R,S*)-ketamine at time points corresponding to 24 and 48 h post CPP conditioning. These results demonstrate the profound effect that (*R,S*)-ketamine has on reward-related behaviors and opens up many avenues including, investigating temporal effects of ketamine treatment at later time points following conditioning, the neurocircuit mechanisms mediating this prolonged ketamine effect on morphine-induced CPP, and the specificity for drug-context associations versus other forms of memory. With the ever increasing use of ketamine as an antidepressant in major depressive disorder (Berman et al., 2000;Diazgranados et al., 2010;Ibrahim et al., 2011;Zarate et al., 2012;Murrough et al., 2013b), applying its therapeutic use to patients suffering from substance use disorders holds potential value as an alternative treatment option.

### Limitations to the use of ketamine as a treatment option for substance use disorders

Despite its therapeutic value, ketamine has undesirable side effects including drowsiness, confusion, dizziness, and dissociative psychiatric side effects (Zarate et al., 2006;Diazgranados et al., 2010;Ibrahim et al., 2011;Murrough et al., 2013a). Additionally, evidence suggests that ketamine impairs cognition and memory (Harris et al., 1975;Ghoneim et al., 1985;Malhotra et al., 1996;Newcomer et al., 1999;Pfenninger et al., 2002;Morgan et al., 2004;Honey et al., 2005;Mathew et al., 2010;Driesen et al., 2013) and may cause urological effects (Middela and Pearce, 2011). A limitation of ketamine use as a treatment option for substance use disorders is its abuse potential (Liu et al., 2016). However, controlled studies in patients addressing the abuse potential of low-dose ketamine are lacking and if the long-lasting ketamine effects shown here in mice translate to human patients, the abuse liability can be mitigated by monthly physician-administered injections.

## Conclusions

Here, we found that morphine conditioning in a three-compartment apparatus that elicits robust CPP was sufficient to evoke anxiety-like behaviors in mice. Additionally, we provided evidence that acute ketamine pretreatment produces anxiolytic-like behaviors and blocks morphine-induced CPP for a prolonged time period, suggesting that ketamine is a potential option for attenuating negative reinforcement as well as learned associations that are implicated in substance use disorders.

## Acknowledgements

We thank Dr. Diane McCloskey for edits and comments on this project as well as Dr. Patrick Randall and Dr. Zheng-Ming Ding for their comments on the project. The study was supported by the Brain & Behavioral Research NARSAD Young Investigator Award (27364NG), the Pennsylvania State Junior Faculty Scholar Award (NG), the Pennsylvania Department of Health using Tobacco CURE Funds (NG), and the Pennsylvania State Research Allocation Project Grant (NG). Morphine was provided by the Drug Supply Program of NIDA NIH.

## Author Contributions Statement

G.M., H.G, H.E.J., D.S.M., S.S. Y.S., and N.M.G. designed the experiments, performed the analyses, and wrote the manuscript. H.E.J., G.M., H.G., D.S.M., and S.S performed behavioral training and testing.

---

**Supplementary Figure 1.**
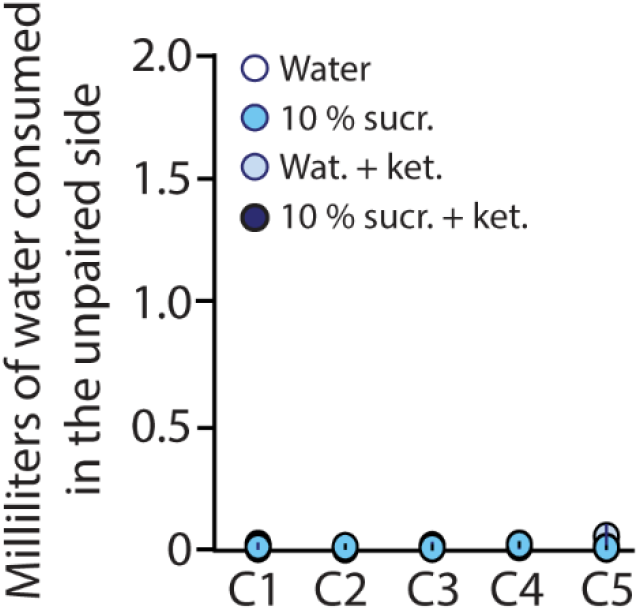
Summary showing that there is no significant difference in the amount of water consumed in the most preferred side among all groups (F_(12, 140)_=0.596, p=0.843, two-way repeated measures ANOVA).

**Supplementary Figure 2.**
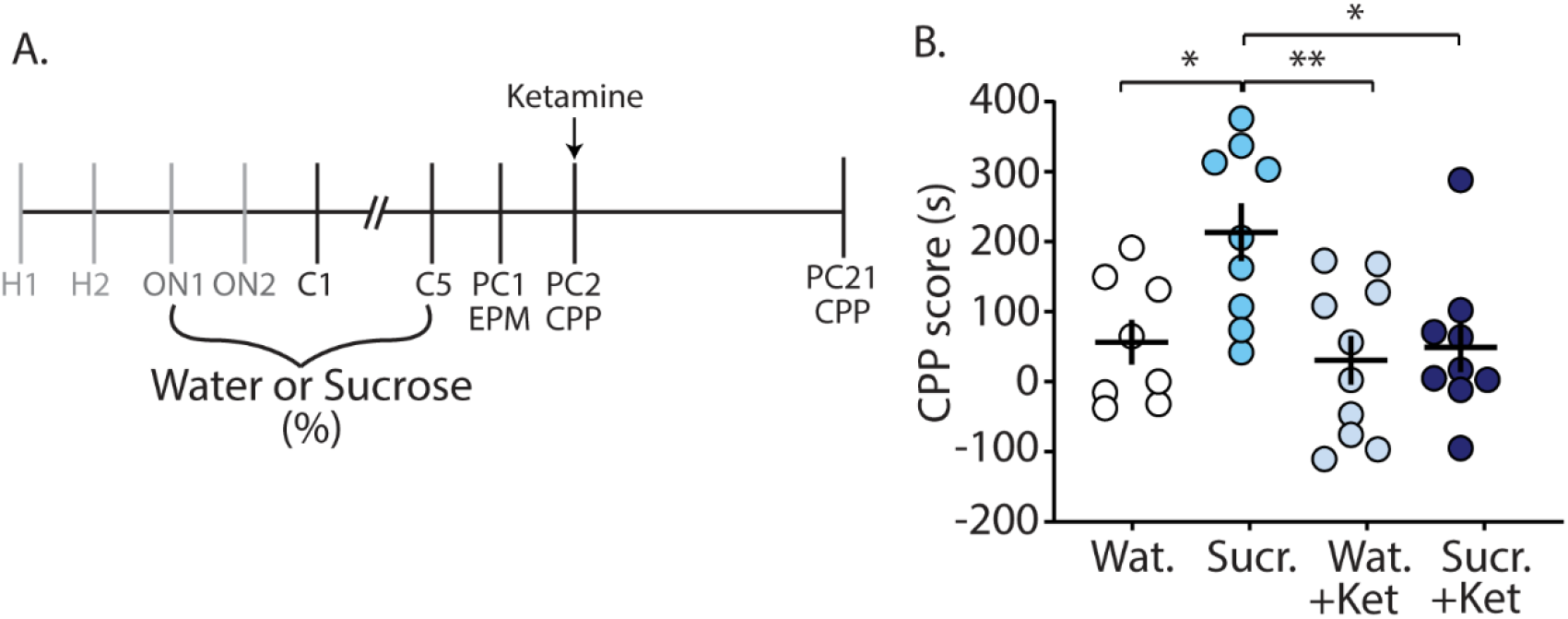
(R,S)-ketamine administration during early abstinence blocks the prolonged retention of sucrose-induced CPP at post conditioning day 21. (A) Time line and drug regimen of the behavioral procedure. (*R,S*)-ketamine (10 mg/kg, i.p.) was injected 30 min. prior to the first CPP test on post conditioning day 2 (PC2). (B) Summary showing that oral self-administration of sucrose produced CPP for the sucrose-paired context 21 days after conditioning. This prolonged expression of sucrose-induced CPP was blocked by (*R,S*)-ketamine when injected 30 min prior to testing on PC2 (F_(3, 32)_=5.51, p=0.004, one-way ANOVA, Bonferroni post hoc test).

## References cited

Akillioglu, K., Melik, E.B., Melik, E., and Boga, A. (2012). Effect of ketamine on exploratory behaviour in BALB/C and C57BL/6 mice. Pharmacology Biochemistry and Behavior 100,513–517.

Aston-Jones, G., Delfs, J.M., Druhan, J., and Zhu, Y. (1999). The bed nucleus of the stria terminalis. A target site for noradrenergic actions in opiate withdrawal. Ann N Y Acad Sci 877,486–498.

Aston-Jones, G., and Harris, G.C. (2004). Brain substrates for increased drug seeking during protracted withdrawal. Neuropharmacology 47, 167–179.

Attarzadeh-Yazdi, G., Karimi, S., Azizi, P., Yazdi-Ravandi, S., Hesam, S., and Haghparast, A. (2013). Inhibitory effects of forced swim stress and corticosterone on the acquisition but not expression of morphine-induced conditioned place preference: involvement of glucocorticoid receptor in the basolateral amygdala. Behavioural brain research 252, 339–346.

Baker, T.B., Piper, M.E., Mccarthy, D.E., Majeskie, M.R., and Fiore, M.C. (2004). Addiction motivation reformulated: an affective processing model of negative reinforcement. Psychol Rev 111, 33–51.

Becker, J.a.J., Kieffer, B.L., and Le Merrer, J. (2017). Differential behavioral and molecular alterations upon protracted abstinence from cocaine versus morphine, nicotine, THC and alcohol. Addict Biol22, 1205–1217.

Beilin, B., Rusabrov, Y., Shapira, Y., Roytblat, L., Greemberg, L., Yardeni, I.Z., and Bessler, H. (2007). Low-dose ketamine affects immune responses in humans during the early postoperative period. Br J Anaesth 99, 522–527.

Benturquia, N., Le Guen, S., Canestrelli, C., Lagente, V., Apiou, G., Roques, B.P., and Noble, F. (2007). Specific blockade of morphine- and cocaine-induced reinforcing effects in conditioned place preference by nitrous oxide in mice. Neuroscience 149, 477–486.

Berman, R.M., Cappiello, A., Anand, A., Oren, D.A., Heninger, G.R., Charney, D.S., and Krystal, J.H. (2000). Antidepressant effects of ketamine in depressed patients. Biol Psychiatry 47, 351–354.

Bilbey, D.L., Salem, H., and Grossman, M.H. (1960). The anatomical basis of the straub phenomenon. Br J Pharmacol Chemother 15, 540–543.

Bohn, L.M., Gainetdinov, R.R., Sotnikova, T.D., Medvedev, I.O., Lefkowitz, R.J., Dykstra, L.A., and Caron, M.G. (2003). Enhanced rewarding properties of morphine, but not cocaine, in beta(arrestin)-2 knock-out mice. J Neurosci 23, 10265–10273.

Cabral, A., Ruggiero, R.N., Nobre, M.J., Brandao, M.L., and Castilho, V.M. (2009). GABA and opioid mechanisms of the central amygdala underlie the withdrawal-potentiated startle from acute morphine. Prog Neuropsychopharmacol Biol Psychiatry 33, 334–344.

Cai, Y.-C., Ma, L., Fan, G.-H., Zhao, J., Jiang, L.-Z., and Pei, G. (1997). Activation of *N*-Methyl-d-Aspartate Receptor Attenuates Acute Responsiveness of δ-Opioid Receptors. Molecular Pharmacology 51, 583–587.

Conklin, C.A., and Perkins, K.A. (2005). Subjective and reinforcing effects of smoking during negative mood induction. J Abnorm Psychol 114, 153–164.

Cooney, N.L., Litt, M.D., Morse, P.A., Bauer, L.O., and Gaupp, L. (1997). Alcohol cue reactivity, negative-mood reactivity, and relapse in treated alcoholic men. J Abnorm Psychol 106, 243–250.

Da Silva, F.C.C., Do Carmo De Oliveira Cito, M., Da Silva, Moura, B.A., De Aquino Neto, M.R., Feitosa, M.L., De Castro Chaves, R., Macedo, D.S., De Vasconcelos, S.M.M., De França Fonteles, M.M., and De Sousa, F.C.F. (2010). Behavioral alterations and pro-oxidant effect of a single ketamine administration to mice. Brain Research Bulletin 83, 9–15.

Dakwar, E., Hart, C.L., Levin, F.R., Nunes, E.V., and Foltin, R.W. (2017). Cocaine self-administration disrupted by the N-methyl-D-aspartate receptor antagonist ketamine: a randomized, crossover trial. Molecular Psychiatry 22, 76–81.

Dawson, G.R., and Tricklebank, M.D. (1995). Use of the elevated plus maze in the search for novel anxiolytic agents. Trends in pharmacological sciences 16, 33–36.

Delfs, J.M., Zhu, Y., Druhan, J.P., and Aston-Jones, G. (2000). Noradrenaline in the ventral forebrain is critical for opiate withdrawal-induced aversion. Nature 403, 430–434.

Diana, M., Pistis, M., Muntoni, A., and Gessa, G. (1995). Profound decrease of mesolimbic dopaminergic neuronal activity in morphine withdrawn rats. J Pharmacol Exp Ther 272, 781–785.

Diazgranados, N., Ibrahim, L., Brutsche, N.E., Newberg, A., Kronstein, P., Khalife, S., Kammerer, W.A., Quezado, Z., Luckenbaugh, D.A., Salvadore, G., Machado-Vieira, R., Manji, H.K., and Zarate, C.A., Jr. (2010). A randomized add-on trial of an N-methyl-D-aspartate antagonist in treatment-resistant bipolar depression. Arch Gen Psychiatry 67, 793–802.

Driesen, N.R., Mccarthy, G., Bhagwagar, Z., Bloch, M.H., Calhoun, V.D., D’souza, D.C., Gueorguieva, R., He, G., Leung, H.C., Ramani, R., Anticevic, A., Suckow, R.F., Morgan, P.T., and Krystal, J.H. (2013). The impact of NMDA receptor blockade on human working memory-related prefrontal function and connectivity. Neuropsychopharmacology 38, 2613–2622.

Ebert, B., Mikkelsen, S., Thorkildsen, C., and Borgbjerg, F.M. (1997). Norketamine, the main metabolite of ketamine, is a non-competitive NMDA receptor antagonist in the rat cortex and spinal cord. Eur J Pharmacol 333, 99–104.

Engin, E., Treit, D., and Dickson, C.T. (2009a). Anxiolytic- and antidepressant-like properties of ketamine in behavioral and neurophysiological animal models. Neuroscience 161, 359–369.

Engin, E., Treit, D., and Dickson, C.T. (2009b). Anxiolytic- and antidepressant-like properties of ketamine in behavioral and neurophysiological animal models. Neuroscience 161, 359–369.

Evans, C.J., and Cahill, C.M. (2016). Neurobiology of opioid dependence in creating addiction vulnerability. F1000Res 5.

Filibeck, U., and Castellano, C. (1980). Strain dependent effects of ketamine on locomotor activity and antinociception in mice. Pharmacol Biochem Behav 13, 443–447.

Fox, H.C., Bergquist, K.L., Hong, K.I., and Sinha, R. (2007). Stress-induced and alcohol cue-induced craving in recently abstinent alcohol-dependent individuals. Alcohol Clin Exp Res 31, 395–403.

Fraga, D.B., Olescowicz, G., Moretti, M., Siteneski, A., Tavares, M.K., Azevedo, D., Colla, A.R.S., and Rodrigues, A.L.S. (2018). Anxiolytic effects of ascorbic acid and ketamine in mice. J Psychiatr Res 100, 16–23.

Freet, C.S., Wheeler, R.A., Leuenberger, E., Mosblech, N.A., and Grigson, P.S. (2013). Fischer rats are more sensitive than Lewis rats to the suppressive effects of morphine and the aversive kappa-opioid agonist spiradoline. Behav Neurosci 127, 763–770.

Fuentealba, J.A., Forray, M.I., and Gysling, K. (2000). Chronic morphine treatment and withdrawal increase extracellular levels of norepinephrine in the rat bed nucleus of the stria terminalis. J Neurochem 75, 741–748.

Gallego, X., Murtra, P., Zamalloa, T., Canals, J.M., Pineda, J., Amador-Arjona, A., Maldonado, R., and Dierssen, M. (2010). Increased opioid dependence in a mouse model of panic disorder. Front Behav Neurosci 3, 60.

Ganguly, S., Panetta, J.C., Roberts, J.K., and Schuetz, E.G. (2018). Ketamine Pharmacokinetics and Pharmacodynamics Are Altered by P-Glycoprotein and Breast Cancer Resistance Protein Efflux Transporters in Mice. Drug Metab Dispos 46, 1014–1022.

Ghoneim, M.M., Hinrichs, J.V., Mewaldt, S.P., and Petersen, R.C. (1985). Ketamine: behavioral effects of subanesthetic doses. J Clin Psychopharmacol 5, 70–77.

Glue, P., Neehoff, S.M., Medlicott, N.J., Gray, A., Kibby, G., and Mcnaughton, N. (2018). Safety and efficacy of maintenance ketamine treatment in patients with treatment-refractory generalised anxiety and social anxiety disorders. J Psychopharmacol 32, 663–667.

Gold, M.S., Redmond, D.E., Jr., and Kleber, H.D. (1978). Clonidine blocks acute opiate-withdrawal symptoms. Lancet 2, 599–602.

Gold, M.S., Redmond, D.E., Jr., and Kleber, H.D. (1979). Noradrenergic hyperactivity in opiate withdrawal supported by clonidine reversal of opiate withdrawal. Am J Psychiatry 136, 100–102.

Goulart, B.K., De Lima, M.N.M., De Farias, C.B., Reolon, G.K., Almeida, V.R., Quevedo, J., Kapczinski, F., Schröder, N., and Roesler, R. (2010). Ketamine impairs recognition memory consolidation and prevents learning-induced increase in hippocampal brain-derived neurotrophic factor levels. Neuroscience 167, 969–973.

Gracy, K.N., Dankiewicz, L.A., and Koob, G.F. (2001). Opiate withdrawal-induced fos immunoreactivity in the rat extended amygdala parallels the development of conditioned place aversion. Neuropsychopharmacology 24, 152–160.

Graziane, N.M., Sun, S., Wright, W.J., Jang, D., Liu, Z., Huang, Y.H., Nestler, E.J., Wang, Y.T., Schluter, O.M., and Dong, Y. (2016). Opposing mechanisms mediate morphine- and cocaine-induced generation of silent synapses. Nat Neurosci 19, 915–925.

Gregus, A., Wintink, A.J., Davis, A.C., and Kalynchuk, L.E. (2005). Effect of repeated corticosterone injections and restraint stress on anxiety and depression-like behavior in male rats. Behav Brain Res 156, 105–114.

Gremel, C.M., Gabriel, K.I., and Cunningham, C.L. (2006). Topiramate does not affect the acquisition or expression of ethanol conditioned place preference in DBA/2J or C57BL/6J mice. Alcohol Clin Exp Res 30, 783–790.

Grisel, J.E., Bartels, J.L., Allen, S.A., and Turgeon, V.L. (2008). Influence of beta-Endorphin on anxious behavior in mice: interaction with EtOH. Psychopharmacology (Berl) 200, 105–115.

Haghparast, A., Fatahi, Z., Alamdary, S.Z., Reisi, Z., and Khodagholi, F. (2014). Changes in the Levels of p-ERK, p-CREB, and c-fos in Rat Mesocorticolimbic Dopaminergic System After Morphine-Induced Conditioned Place Preference: The Role of Acute and Subchronic Stress. Cellular and Molecular Neurobiology 34, 277–288.

Handley, S.L., and Mcblane, J.W. (1993). An assessment of the elevated X-maze for studying anxiety and anxiety-modulating drugs. Journal of pharmacological and toxicological methods 29, 129–138.

Hanse, E., Seth, H., and Riebe, I. (2013). AMPA-silent synapses in brain development and pathology. Nat Rev Neurosci 14, 839–850.

Harris, J.A., Biersner, R.J., Edwards, D., and Bailey, L.W. (1975). Attention, learning, and personality during ketamine emergence: a pilot study. Anesth Analg 54, 169–172.

Hearing, M., Graziane, N., Dong, Y., and Thomas, M.J. (2018). Opioid and Psychostimulant Plasticity: Targeting Overlap in Nucleus Accumbens Glutamate Signaling. Trends Pharmacol Sci 39, 276–294.

Heinrichs, S.C., Menzaghi, F., Schulteis, G., Koob, G.F., and Stinus, L. (1995). Suppression of corticotropin-releasing factor in the amygdala attenuates aversive consequences of morphine withdrawal. Behav Pharmacol 6, 74–80.

Honey, G.D., Honey, R.A., Sharar, S.R., Turner, D.C., Pomarol-Clotet, E., Kumaran, D., Simons, J.S., Hu, X., Rugg, M.D., Bullmore, E.T., and Fletcher, P.C. (2005). Impairment of specific episodic memory processes by sub-psychotic doses of ketamine: the effects of levels of processing at encoding and of the subsequent retrieval task. Psychopharmacology(Berl) 181, 445–457.

Huston, J.P., Silva, M.A., Topic, B., and Müller, C.P. (2013). What’s conditioned in conditioned place preference? Trends Pharmacol Sci 34, 162–166.

Huys, Q.J.M., Tobler, P.N., Hasler, G., and Flagel, S.B. (2014). “Chapter 3 - The role of learning-related dopamine signals in addiction vulnerability,” in Progress in Brain Research, eds. M. Diana, G. Di Chiara & P. Spano. Elsevier), 31–77.

Ibrahim, L., Diazgranados, N., Luckenbaugh, D.A., Machado-Vieira, R., Baumann, J., Mallinger, A.G., and Zarate, C.A., Jr. (2011). Rapid decrease in depressive symptoms with an N-methyl-d-aspartate antagonist in ECT-resistant major depression. Prog Neuropsychopharmacol Biol Psychiatry 35, 1155–1159.

Ivan Ezquerra-Romano, I., Lawn, W., Krupitsky, E., and Morgan, C.J.A. (2018). Ketamine for the treatment of addiction: Evidence and potential mechanisms. Neuropharmacology 142, 72–82.

Jansen, K. (1989). Near death experience and the NMDA receptor. BMJ (Clinical research ed.) 298, 1708–1708.

Jones, J.L., Mateus, C.F., Malcolm, R.J., Brady, K.T., and Back, S.E. (2018). Efficacy of Ketamine in the Treatment of Substance Use Disorders: A Systematic Review. Frontiers in psychiatry 9, 277–277.

Jonkman, K., Dahan, A., Van De Donk, T., Aarts, L., Niesters, M., and Van Velzen, M. (2017). Ketamine for pain. F1000Research 6, F1000 Faculty Rev-1711.

Karimi, S., Attarzadeh-Yazdi, G., Yazdi-Ravandi, S., Hesam, S., Azizi, P., Razavi, Y., and Haghparast, A. (2014). Forced swim stress but not exogenous corticosterone could induce the reinstatement of extinguished morphine conditioned place preference in rats: involvement of glucocorticoid receptors in the basolateral amygdala. Behav Brain Res 264, 43–50.

Khorramzadeh, E., and Lotfy, A.O. (1973). The use of ketamine in psychiatry. Psychosomatics 14, 344–346.

Kohrs, R., and Durieux, M.E. (1998). Ketamine: teaching an old drug new tricks. Anesth Analg 87, 1186–1193.

Koks, S., Soosaar, A., Voikar, V., Bourin, M., and Vasar, E. (1999). BOC-CCK-4, CCK(B)receptor agonist, antagonizes anxiolytic-like action of morphine in elevated plus-maze. Neuropeptides 33, 63–69.

Kolp, E., Friedman, H.L., Young, M.S., and Krupitsky, E. (2006). Ketamine Enhanced Psychotherapy: Preliminary Clinical Observations on Its Effectiveness in Treating Alcoholism. The Humanistic Psychologist 34, 399–422.

Koo, J.W., Lobo, M.K., Chaudhury, D., Labonte, B., Friedman, A., Heller, E., Pena, C.J., Han, M.H., and Nestler, E.J. (2014). Loss of BDNF signaling in D1R-expressing NAc neurons enhances morphine reward by reducing GABA inhibition. Neuropsychopharmacology 39, 2646–2653.

Koob, G.F., and Le Moal, M. (2008). Review. Neurobiological mechanisms for opponent motivational processes in addiction. Philos Trans R Soc Lond B Biol Sci363, 3113–3123.

Krupitsky, E., Burakov, A., Romanova, T., Dunaevsky, I., Strassman, R., and Grinenko, A. (2002a). Ketamine psychotherapy for heroin addiction: immediate effects and two-year follow-up. J Subst Abuse Treat 23, 273–283.

Krupitsky, E., Burakov, A., Romanova, T., Dunaevsky, I., Strassman, R., and Grinenko, A. (2002b). Ketamine psychotherapy for heroin addiction: immediate effects and two-year follow-up. Journal of Substance Abuse Treatment 23, 273–283.

Krupitsky, E.M., Burakov, A.M., Dunaevsky, I.V., Romanova, T.N., Slavina, T.Y., and Grinenko, A.Y. (2007). Single Versus Repeated Sessions of Ketamine-Assisted Psychotherapy for People with Heroin Dependence. Journal of Psychoactive Drugs 39, 13–19.

Krupitsky, E.M., and Grinenko, A.Y. (1997). Ketamine Psychedelic Therapy (KPT): A Review of the Results of Ten Years of Research. Journal of Psychoactive Drugs 29, 165–183.

Krystal, J.H., Karper, L.P., Seibyl, J.P., Freeman, G.K., Delaney, R., Bremner, J.D., Heninger, G.R., Bowers, M.B., Jr., and Charney, D.S. (1994). Subanesthetic effects of the noncompetitive NMDA antagonist, ketamine, in humans. Psychotomimetic, perceptual, cognitive, and neuroendocrine responses. Arch Gen Psychiatry 51, 199–214.

Kubota, T., Hirota, K., Yoshida, H., Takahashi, S., Anzawa, N., Ohkawa, H., Kushikata, T., and Matsuki, A. (1999). Effects of sedatives on noradrenaline release from the medial prefrontal cortex in rats. Psychopharmacology (Berl) 146, 335–338.

Laskowski, K., Stirling, A., Mckay, W.P., and Lim, H.J. (2011). A systematic review of intravenous ketamine for postoperative analgesia. Can J Anaesth 58, 911–923.

Li, S.X., Shi, J., Epstein, D.H., Wang, X., Zhang, X.L., Bao, Y.P., Zhang, D., Zhang, X.Y., Kosten, T.R., and Lu, L. (2009). Circadian alteration in neurobiology during 30 days of abstinence in heroin users. Biol Psychiatry 65, 905–912.

Li, X., Martinez-Lozano Sinues, P., Dallmann, R., Bregy, L., Hollmen, M., Proulx, S., Brown, S.A., Detmar, M., Kohler, M., and Zenobi, R. (2015). Drug Pharmacokinetics Determined by Real-Time Analysis of Mouse Breath. Angew Chem Int Ed Engl 54, 7815–7818.

Lindholm, J.S.O., Autio, H., Vesa, L., Antila, H., Lindemann, L., Hoener, M.C., Skolnick, P., Rantamäki, T., and Castrén, E. (2012). The antidepressant-like effects of glutamatergic drugs ketamine and AMPA receptor potentiator LY 451646 are preserved in bdnf+/-heterozygous null mice. Neuropharmacology 62, 391–397.

Liu, Y., Lin, D., Wu, B., and Zhou, W. (2016). Ketamine abuse potential and use disorder. Brain Research Bulletin 126, 68–73.

Lodge, D., Anis, N.A., and Burton, N.R. (1982). Effects of optical isomers of ketamine on excitation of cat and rat spinal neurones by amino acids and acetylcholine. Neurosci Lett 29, 281–286.

Loix, S., De Kock, M., and Henin, P. (2011). The anti-inflammatory effects of ketamine: state of the art. Acta Anaesthesiol Belg62, 47–58.

Louderback, K.M., Wills, T.A., Muglia, L.J., and Winder, D.G. (2013). Knockdown of BNST GluN2B-containing NMDA receptors mimics the actions of ketamine on novelty-induced hypophagia. Transl Psychiatry 3, e331.

Lu, L., Chen, H., Su, W., Ge, X., Yue, W., Su, F., and Ma, L. (2005). Role of withdrawal in reinstatement of morphine-conditioned place preference. Psychopharmacology 181, 90–100.

Lydic, R., and Baghdoyan, H.A. (2002). Ketamine and MK-801 decrease acetylcholine release in the pontine reticular formation, slow breathing, and disrupt sleep. Sleep 25, 617–622.

Maj, M., Turchan, J., Śmiałowska, M., and Przewłocka, B. (2003). Morphine and cocaine influence on CRF biosynthesis in the rat central nucleus of amygdala. Neuropeptides 37, 105–110.

Malhotra, A.K., Pinals, D.A., Weingartner, H., Sirocco, K., Missar, C.D., Pickar, D., and Breier, A. (1996). NMDA receptor function and human cognition: the effects of ketamine in healthy volunteers. Neuropsychopharmacology 14, 301–307.

Martins, S.S., Fenton, M.C., Keyes, K.M., Blanco, C., Zhu, H., and Storr, C.L. (2012). Mood and anxiety disorders and their association with non-medical prescription opioid use and prescription opioid-use disorder: longitudinal evidence from the National Epidemiologic Study on Alcohol and Related Conditions. Psychol Med 42, 1261–1272.

Mathew, S.J., Murrough, J.W., Aan Het Rot, M., Collins, K.A., Reich, D.L., and Charney, D.S. (2010). Riluzole for relapse prevention following intravenous ketamine in treatment-resistant depression: a pilot randomized, placebo-controlled continuation trial. Int J Neuropsychopharmacol 13, 71–82.

Maxwell, C.R., Ehrlichman, R.S., Liang, Y., Trief, D., Kanes, S.J., Karp, J., and Siegel, S.J. (2006). Ketamine produces lasting disruptions in encoding of sensory stimuli. J Pharmacol Exp Ther 316, 315–324.

Mcdevitt, D.S., and Graziane, N.M. (2018). Neuronal mechanisms mediating pathological reward-related behaviors: A focus on silent synapses in the nucleus accumbens. Pharmacol Res 136, 90–96.

Mcdevitt, D.S., and Graziane, N.M. (2019). Timing of Morphine Administration Differentially Alters Paraventricular Thalamic Neuron Activity. eNeuro 6, ENEURO.0377-0319.2019.

Middela, S., and Pearce, I. (2011). Ketamine-induced vesicopathy: a literature review. Int J Clin Pract 65, 27–30.

Miller, D.B., Dougherty, J.A., and Wikler, A. (1979). Interoceptive conditioning through repeated suppression of morphine-abstinence. II. Relapse-testing. The Pavlovian journal of biological science 14, 170–176.

Moaddel, R., Abdrakhmanova, G., Kozak, J., Jozwiak, K., Toll, L., Jimenez, L., Rosenberg, A., Tran, T., Xiao, Y., Zarate, C.A., and Wainer, I.W. (2013). Sub-anesthetic concentrations of (R,S)-ketamine metabolites inhibit acetylcholine-evoked currents in alpha7 nicotinic acetylcholine receptors. Eur J Pharmacol 698, 228–234.

Morgan, C.J., Mofeez, A., Brandner, B., Bromley, L., and Curran, H.V. (2004). Acute effects of ketamine on memory systems and psychotic symptoms in healthy volunteers. Neuropsychopharmacology29, 208–218.

Murrough, J.W., Iosifescu, D.V., Chang, L.C., Al Jurdi, R.K., Green, C.E., Perez, A.M., Iqbal, S., Pillemer, S., Foulkes, A., Shah, A., Charney, D.S., and Mathew, S.J. (2013a). Antidepressant efficacy of ketamine in treatment-resistant major depression: a two-site randomized controlled trial. Am J Psychiatry 170, 1134–1142.

Murrough, J.W., Perez, A.M., Pillemer, S., Stern, J., Parides, M.K., Aan Het Rot, M., Collins, K.A., Mathew, S.J., Charney, D.S., and Iosifescu, D.V. (2013b). Rapid and longer-term antidepressant effects of repeated ketamine infusions in treatment-resistant major depression. Biol Psychiatry 74, 250–256.

Newcomb, M.D., and Bentler, P.M. (1988). Impact of adolescent drug use and social support on problems of young adults: A longitudinal study. Journal of Abnormal Psychology 97, 64–75.

Newcomer, J.W., Farber, N.B., Jevtovic-Todorovic, V., Selke, G., Melson, A.K., Hershey, T., Craft, S., and Olney, J.W. (1999). Ketamine-induced NMDA receptor hypofunction as a model of memory impairment and psychosis. Neuropsychopharmacology 20, 106–118.

O’brien, C.P. (1975). Experimental analysis of conditioning factors in human narcotic addiction. Pharmacol Rev 27, 533–543.

O’brien, C.P., Childress, A.R., Mclellan, A.T., and Ehrman, R. (1992). Classical conditioning in drug-dependent humans. Ann N Y Acad Sci 654, 400–415.

O’brien Cp, E.R., Ternes Jw (1986). “Classical conditioning in human opioid dependence.,” in Behavioral analysis of drug dependence, ed. S.I. Goldberg S. (Orlando, FL:: Academic), 329–356.

Pellow, S., Chopin, P., File, S.E., and Briley, M. (1985a). Validation of open:closed arm entries in an elevated plus-maze as a measure of anxiety in the rat. J Neurosci Methods 14, 149–167.

Pellow, S., Chopin, P., File, S.E., and Briley, M. (1985b). Validation of open:closed arm entries in an elevated plus-maze as a measure of anxiety in the rat. Journal of neuroscience methods 14, 149–167.

Perkins, K.A., and Grobe, J.E. (1992). Increased desire to smoke during acute stress. Br J Addict 87, 1037–1040.

Pfenninger, E.G., Durieux, M.E., and Himmelseher, S. (2002). Cognitive impairment after small-dose ketamine isomers in comparison to equianalgesic racemic ketamine in human volunteers. Anesthesiology 96, 357–366.

Ribeiro Do Couto, B., Aguilar, M.A., Manzanedo, C., Rodríguez-Arias, M., Armario, A., and Miñarro, J. (2006). Social stress is as effective as physical stress in reinstating morphine-induced place preference in mice. Psychopharmacology 185, 459–470.

Robinson, T.E., and Kolb, B. (1999). Morphine alters the structure of neurons in the nucleus accumbens and neocortex of rats. Synapse 33, 160–162.

Roytblat, L., Talmor, D., Rachinsky, M., Greemberg, L., Pekar, A., Appelbaum, A., Gurman, G.M., Shapira, Y., and Duvdenani, A. (1998). Ketamine attenuates the interleukin-6 response after cardiopulmonary bypass. Anesth Analg 87, 266–271.

Sabino, V., Narayan, A.R., Zeric, T., Steardo, L., and Cottone, P. (2013). mTOR activation is required for the anti-alcohol effect of ketamine, but not memantine, in alcohol-preferring rats. Behavioural brain research 247, 9–16.

Sasaki, K., Fan, L.W., Tien, L.T., Ma, T., Loh, H.H., and Ho, I.K. (2002). The interaction of morphine and gamma-aminobutyric acid (GABA)ergic systems in anxiolytic behavior: using mu-opioid receptor knockout mice. Brain Res Bull57, 689–694.

Scheller, M., Bufler, J., Hertle, I., Schneck, H.J., Franke, C., and Kochs, E. (1996). Ketamine blocks currents through mammalian nicotinic acetylcholine receptor channels by interaction with both the open and the closed state. Anesth Analg 83, 830–836.

Shadli, S.M., Kawe, T., Martin, D., Mcnaughton, N., Neehoff, S., and Glue, P. (2018). Ketamine Effects on EEG during Therapy of Treatment-Resistant Generalized Anxiety and Social Anxiety. Int J Neuropsychopharmacol.

Shaham, Y., Rajabi, H., and Stewart, J. (1996). Relapse to heroin-seeking in rats under opioid maintenance: the effects of stress, heroin priming, and withdrawal. The Journal of neuroscience: the official journal of the Society for Neuroscience 16, 1957–1963.

Shi, J., Li, S.X., Zhang, X.L., Wang, X., Le Foll, B., Zhang, X.Y., Kosten, T.R., and Lu, L. (2009). Time-dependent neuroendocrine alterations and drug craving during the first month of abstinence in heroin addicts. Am J Drug Alcohol Abuse 35, 267–272.

Shin, I.C., Kim, H.C., Swanson, J., Hong, J.T., and Oh, K.W. (2003). Anxiolytic effects of acute morphine can be modulated by nitric oxide systems. Pharmacology 68, 183–189.

Silvestre, J.S., Nadal, R., Pallares, M., and Ferre, N. (1997). Acute effects of ketamine in the holeboard, the elevated-plus maze, and the social interaction test in Wistar rats. Depress Anxiety 5, 29–33.

Sinha, R. (2008). Chronic stress, drug use, and vulnerability to addiction. Annals of the New York Academy of Sciences 1141, 105–130.

Smith, R.J., and Aston-Jones, G. (2008). Noradrenergic transmission in the extended amygdala: role in increased drug-seeking and relapse during protracted drug abstinence. Brain Struct Funct 213, 43–61.

Sofia, R.D., and Harakal, J.J. (1975). Evaluation of ketamine HCl for anti-depressant activity. Arch Int Pharmacodyn Ther 214, 68–74.

Solomon, R.L., and Corbit, J.D. (1978). An Opponent-Process Theory of Motivation. The American Economic Review 68, 12–24.

Suzuki, T., Kato, H., Aoki, T., Tsuda, M., Narita, M., and Misawa, M. (2000). Effects of the non-competitive NMDA receptor antagonist ketamine on morphine-induced place preference in mice. Life Sci 67, 383–389.

Taylor, J.H., Landeros-Weisenberger, A., Coughlin, C., Mulqueen, J., Johnson, J.A., Gabriel, D., Reed, M.O., Jakubovski, E., and Bloch, M.H. (2018). Ketamine for Social Anxiety Disorder: A Randomized, Placebo-Controlled Crossover Trial. Neuropsychopharmacology 43, 325–333.

Tyler, M.W., Yourish, H.B., Ionescu, D.F., and Haggarty, S.J. (2017). Classics in Chemical Neuroscience: Ketamine. ACS Chemical Neuroscience 8, 1122–1134.

Tzschentke, T.M. (2007). Measuring reward with the conditioned place preference (CPP) paradigm: update of the last decade. Addict Biol 12, 227–462.

Wang, J., Fang, Q., Liu, Z., and Lu, L. (2006). Region-specific effects of brain corticotropin-releasing factor receptor type 1 blockade on footshock-stress-or drug-priming-induced reinstatement of morphine conditioned place preference in rats. Psychopharmacology 185, 19–28.

Wang, M., Wong, A.H., and Liu, F. (2012). Interactions between NMDA and dopamine receptors: A potential therapeutic target. Brain Research 1476, 154–163.

Weisman, H. (1971). Anesthesia for pediatric ophthalmology. Ann Ophthalmol3, 229–232.

Wetter, D.W., Smith, S.S., Kenford, S.L., Jorenby, D.E., Fiore, M.C., Hurt, R.D., Offord, K.P., and Baker, T.B. (1994). Smoking outcome expectancies: factor structure, predictive validity, and discriminant validity. J Abnorm Psychol 103, 801–811.

Whitaker, Leslie r., Degoulet, M., and Morikawa, H. (2013). Social Deprivation Enhances VTA Synaptic Plasticity and Drug-Induced Contextual Learning. Neuron 77, 335–345.

Wikler, A. (2013). Opioid Dependence: Mechanisms and Treatment. Springer US.

Xue, Y.X., Luo, Y.X., Wu, P., Shi, H.S., Xue, L.F., Chen, C., Zhu, W.L., Ding, Z.B., Bao, Y.P., Shi, J., Epstein, D.H., Shaham, Y., and Lu, L. (2012). A memory retrieval-extinction procedure to prevent drug craving and relapse. Science 336, 241–245.

Zanos, P., Moaddel, R., Morris, P.J., Georgiou, P., Fischell, J., Elmer, G.I., Alkondon, M., Yuan, P., Pribut, H.J., Singh, N.S., Dossou, K.S.S., Fang, Y., Huang, X.-P., Mayo, C.L., Wainer, I.W., Albuquerque, E.X., Thompson, S.M., Thomas, C.J., Zarate Jr, C.A., and Gould, T.D. (2016). NMDAR inhibition-independent antidepressant actions of ketamine metabolites. Nature 533, 481.

Zanos, P., Moaddel, R., Morris, P.J., Riggs, L.M., Highland, J.N., Georgiou, P., Pereira, E.F.R., Albuquerque, E.X., Thomas, C.J., Zarate, C.A., Jr., and Gould, T.D. (2018). Ketamine and Ketamine Metabolite Pharmacology: Insights into Therapeutic Mechanisms. Pharmacological reviews 70, 621–660.

Zarate, C.A., Jr., Brutsche, N., Laje, G., Luckenbaugh, D.A., Venkata, S.L., Ramamoorthy, A., Moaddel, R., and Wainer, I.W. (2012). Relationship of ketamine’s plasma metabolites with response, diagnosis, and side effects in major depression. Biol Psychiatry 72, 331–338.

Zarate, C.A., Jr., Singh, J.B., Carlson, P.J., Brutsche, N.E., Ameli, R., Luckenbaugh, D.A., Charney, D.S., and Manji, H.K. (2006). A randomized trial of an N-methyl-D-aspartate antagonist in treatment-resistant major depression. Arch Gen Psychiatry 63, 856–864.

Zhai, H., Wu, P., Chen, S., Li, F., Liu, Y., and Lu, L. (2008). Effects of scopolamine and ketamine on reconsolidation of morphine conditioned place preference in rats. Behav Pharmacol 19, 211–216.

Zhang, L.-M., Zhou, W.-W., Ji, Y.-J., Li, Y., Zhao, N., Chen, H.-X., Xue, R., Mei, X.-G., Zhang, Y.-Z., Wang, H.-L., and Li, Y.-F. (2015). Anxiolytic effects of ketamine in animal models of posttraumatic stress disorder. Psychopharmacology 232, 663–672.

Zinser, M.C., Baker, T.B., Sherman, J.E., and Cannon, D.S. (1992). Relation between self-reported affect and drug urges and cravings in continuing and withdrawing smokers. J Abnorm Psychol 101, 617–629.

